# Gaining comprehensive biological insight into chloroplast RNA editing by performing a broad-spectrum RNA-seq analysis

**DOI:** 10.1101/2020.06.02.129577

**Authors:** Aidi Zhang, Jing Fang, Xiaohan Jiang, Tengfei Wang, Xiujun Zhang

**Author notes:** Correspondence, Wuhan Botanical Garden, Chinese Academy of Sciences, Wuhan 430072, China.

## Abstract

**Background:** RNA editing is a post-transcriptional modification that complement variation at the DNA level. Until now, different RNA editing systems were found in the major eukaryotic lineages. However, the evolution trajectory in plant chloroplast remains unclear. To gain a better understanding of RNA editing in plant chloroplast, in this study, based on publicly available RNA-seq data across three plant lineages (fern, gymnosperm, and angiosperm), we provided a detailed analysis of RNA editing events in plant chloroplasts and discussed the evolution of RNA editing in land plants.

**Results:** There were a total of 5,389 editing sites located in leaf chloroplast identified across 21 plants after rigorous screening. We found that the cluster of RNA editing sites across 21 plants complied with the phylogenetic tree based on linked protein sequences approximately, and majority (∼ 95%) of the editing events resulted in non-synonymous codon changes, RNA editing occurred in second codon position was mainly the largest. Additionally, RNA editing caused an overall increase in hydrophobicity of the resulting proteins. The analyses also revealed that there is an uneven distribution of editing sites among species, genes, and codon positions, the average RNA editing extent varied among different plant species as well as genes. Finally, we found that the loss of editing sites along angiosperm evolution is mainly occurring by reduce of cytosines content, fern plants has the highest cytosine content, with the evolution of plants, cytosine were lost in RNA edited genes.

**Conclusions:** Many of the identified sites in our study have not been previously reported and represent a valuable data set for future research community. Our findings provide valuable information for evolution of RNA editing in plants.

## Background

RNA editing is a post-transcription process through which the nucleotide specified in the genome template is modified to produce a different transcript, thus contribute to proteomic sequence variation and provide another mechanism for modulating gene expression [1, 2]. In the plant kingdom, RNA editing was first documented over a decade ago in the mitochondria of flowering plants [3, 4], and reported in angiosperm chloroplasts two years later [5], no editing seems to occur in nuclear genome-encoded transcripts. There are two types of RNA editing in plants, the most common type is C-to-U conversion, and the infrequent type is U-to-C conversion that reported only in ferns, mosses, and Lycopodiaceae. RNA editing predominantly take place at the first or second positions of codons, thereby affects the translated regions of protein coding transcripts, occasionally, it can produce a new protein with a different function from the genome-encoded transcript. The amino acids specified by the codons generated by editing in the mRNA are generally better conserved in evolution than the amino acids encoded by the genomic DNA, suggesting that most RNA editing events can restore the evolutionarily conserved amino acid residues in mRNAs. RNA editing thereby is an essential process to maintain essential functions of encoded proteins at the RNA level, for example, pigment deficiency in nightshade/tobacco cybrids is caused by the failure of editing the plastid *ATPase alpha-subunit mRNA* [6].

Numerous studies have demonstrated that RNA editing plays an important role in various plant fundamental processes, such as organelle biogenesis, adaptation to environmental changes, and signal transduction [7-9]. A number of factors are involved in plant RNA editing, and considered to interact with one another to form a large protein complex, termed as editsome, PLS subfamily members of pentatrico peptide repeat (PPR) proteins function in site recognition of the target C, multiple organelle RNA editing factors (MORF) family members are also required for RNA editing at multiple editing sites and are components of the RNA editosome in plants [10, 11]. Almost all the PPR proteins are localized in either chloroplasts or mitochondria where those proteins participate in different facets of RNA metabolism such as RNA splicing, RNA stability, and translational initiation [12]. Despite progress in increasing editing sites identified and the mechanism underlying target recognition, the biological function of RNA editing in plants remains the fundamental question.

More and more recent studies demonstrated that RNA editing is a widespread phenomenon that occurred in the land plants, including the liverworts, mosses, hornworts, lycopods, ferns and flowering plants. However, no instance of RNA editing has yet been reported in *Marchantiidae* and algae, suggesting that RNA editing may have evolved only after the plants established themselves on the land, [13]. Excellent studies in different plants have recently appeared and describe their mechanistic and functional aspects [14-16]. The frequency of organellar RNA editing varies from zero to thousands of sites across plant kingdom, among land plants, early-diverging lineages show the highest numbers of editing sites compared with higher plants that undergoing an extensive loss of editing sites through the substitution of genomic editable cytidines to thymidines [17]. The evidence gathered to date suggests that RNA editing in plant organelles evolved independently from RNA processes in distant evolutionary lineages of animals, fungi, and protozoans when the first plants moved from the aquatic environment onto the land. Evolutionary studies can help to understand the puzzling nature of RNA editing in plants.

The straightforward way to detect RNA editing sites is to compare RNAs with their corresponding DNA templates, sanger sequencing of cDNAs was widely used in the last two decades, though it is time-consuming and prone to underestimate editing sites. In recent years, the advent of next-generation sequencing technologies and the availability of large quantities of RNA sequencing data makes it possible to identify RNA editing sites and quantify their editing extent on a large scale. This strategy allows a transcriptome-wide fast detection of editing sites and has enormous potential to deepen our knowledge of transcriptional processes in plant. Indeed, the number of complete plant organellar genomes and related transcriptome data have considerably grown in the last decade, hence, tens of thousands of editing sites have been identified in more and more plants [18, 19]. However, this strategy is also a challenging task due to its accuracy of mapping the RNA-seq reads against genomic sequence, hence, different bioinformatic strategies have been introduced to improve the detection of RNA editing sites [20-22].

To gain a better understanding of RNA editing in plant chloroplast, in this study, we investigated RNA editing located in chloroplast genome across diverse plants that distributed in three plant lineages: fern, gymnosperm, and angiosperm. Through systematically exploiting enormous RNA-seq data derived from public database, thousands of RNA editing sites were detected, with a lot of these sites have not been previously reported and represent a valuable data set for the research community. Additionally, we provided a detailed analysis of data in land plant chloroplasts and propose a model for the evolution of RNA editing in land plants.

## Results

### Identification of RNA editing sites

To identify RNA editing sites, to represent divergent lineages across plant phylogeny, a series of plant species were selected judging by their availability of transcriptomic data in the SRA database at NCBI and chloroplast genome sequence. Hence, 21 plant species, consisting of 6 ferns, 4 gymnosperms and 11 angiosperms, and corresponding 317 SRA accessions were chosen finally, detailed information of SRA data and plant chloroplast genome accessions were listed in Additional file 1: Table S1. Based on results of RNA-seq data mapping and SNP calling, an automated bioinformatics pipeline implemented in REDO tools was conducted under default thresholds. Consequently, there were a total of 6,011 raw editing sites located in leaf chloroplast genome were detected. However, sequence mismatches that accord with RNA editing occasionally appear in spite of multiple stringent filters, hence, these false positive mismatches should be eliminated. To address this issue, we manually examined all mismatches, only C-to-U and U-to-C editing types were kept, therefore, a total of 5,389 RNA editing sites, consisting of 264 U-to-C and 5,125 C-to-U edit sites, were finally screened out for further analysis.

For the reason that RNA-seq data volume varies among species, several species, such as *Adiantum aleuticum, Histiopteris incisa*, lacked enough RNA-seq data, caused the read depth of these two species was lower than that of other species’, which also reflected the differential accumulation of chloroplast RNA. Furthermore, the read density also varied widely between different genomic regions, ranges from less than 30 to more than 800 in a few species, demonstrating various expression level of genes. The average depth distribution across 21 species was shown in Additional file 6: Figure S1, compared with angiosperms, ferns and gymnosperms have more lower depth, indicating the actual number of RNA editing in ferns and gymnosperms might be underestimated. To evaluate the reliability of editing sites number, PREPACT webserver was also used to predict editing sites, we found that the distribution of predicted number of RNA editing based on PREPACT accorded with that of our prediction based on RNA-seq data basically (see Additional file 6: Figure S2), reflecting our pipeline offered a high performance with reliable results. To describe the attributes of RNA editing sites, we illustrated by one example of samples of *Adiantum aleuticum* (see Additional file 6: Figure S3), which depicts the reliability of RNA editing sites in each species by REDO tools statistically.

### Characteristics of the statistics for RNA editing sites

The statistics of raw results showed that C-to-U was the dominate editing type (nearly ∼95.1%), the next is U-to-C type, other mismatches types were rare (see Figure 1c). After manual inspection of raw results and elimination of mismatches, filtered RNA editing sites across 21 plant species were screened out, detailed corresponding annotation information files were produced simultaneously, as listed in Additional file 2: Table S2, the statistical result was also listed in Table 1, Additional file 3: Table S3 and Figure 1. In terms of codon position, 21.97%, 67.16%, and 10.58% of sites were edited at first, second, and third codon positions respectively (Figure 1b). Furthermore, the statistics of editing types showed that the majority (∼ 95%) of the editing events resulted in non-synonymous codon changes, and the amino acids changes tend to be hydrophobic, the change from hydrophilic to hydrophobic was the highest, followed by the change from hydrophobic to hydrophobic. The most amino acid changes type were Ser-to-Leu and Pro-to-Leu, Serine is hydrophilic, whereas Leucine and Proline are both hydrophobic, as shown in Figure 1d. The above results are in good agreement with previous studies [9], which demonstrated that the RNA editing causes an overall increase in hydrophobicity of the resulting proteins.

**Table 1.** The statistical result of RNA editing sites across 21 plant species

|  | Species | Genome | Raw | Filtered | Percent of edited Genes | Depth |
| --- | --- | --- | --- | --- | --- | --- |
| **Fern** | *Pteris vittata* | MH500228 | 349 | 324 | 0.74 | 102 |
|  | *Adiantum aleuticum* | MH173079 | 422 | 380 | 0.82 | 23 |
|  | *Selaginella moellendorffii* | HM173080 | 1316 | 1316 | 0.96 | 73 |
|  | *Histiopteris incisa* | NC_040220 | 588 | 466 | 0.85 | 27 |
|  | *Cibotium barometz* | NC_037893 | 562 | 549 | 0.85 | 32 |
|  | *Cyrtomium fortunei* | NC_037510 | 778 | 610 | 0.91 | 40 |
| **Gymn** | *Ginkgo biloba* | NC_016986 | 301 | 291 | 0.81 | 228 |
|  | *Picea abies* | HF937082 | 123 | 121 | 0.61 | 450 |
|  | *Cycas revoluta* | NC_020319 | 173 | 140 | 0.44 | 27 |
|  | *Pinus massoniana* | MF564195 | 96 | 93 | 0.59 | 182 |
| **Angi** | *Liriodendron tulipifera* | NC_008326 | 320 | 265 | 0.75 | 113 |
|  | *Nelumbo nucifera* | NC_025339 | 154 | 134 | 0.63 | 295 |
|  | *Nicotiana tabacum* | Z00044 | 107 | 88 | 0.23 | 134 |
|  | *Glycine max* | NC_007942 | 64 | 59 | 0.34 | 195 |
|  | *Populus tremula* | KP861984 | 93 | 89 | 0.43 | 244 |
|  | *Arabidopsis thaliana* | KX551970 | 50 | 45 | 0.24 | 389 |
|  | *Gossypium hirsutum* | NC_007944 | 127 | 100 | 0.37 | 171 |
|  | *Helianthus annuus* | NC_007977 | 59 | 57 | 0.29 | 148 |
|  | *Phoenix dactylifera* | NC_013991 | 121 | 118 | 0.42 | 781 |
|  | *Zea mays* | NC_001666 | 105 | 79 | 0.37 | 814 |
|  | *Oryza sativa* | NC_001320 | 103 | 65 | 0.26 | 550 |

**Fig. 1.**
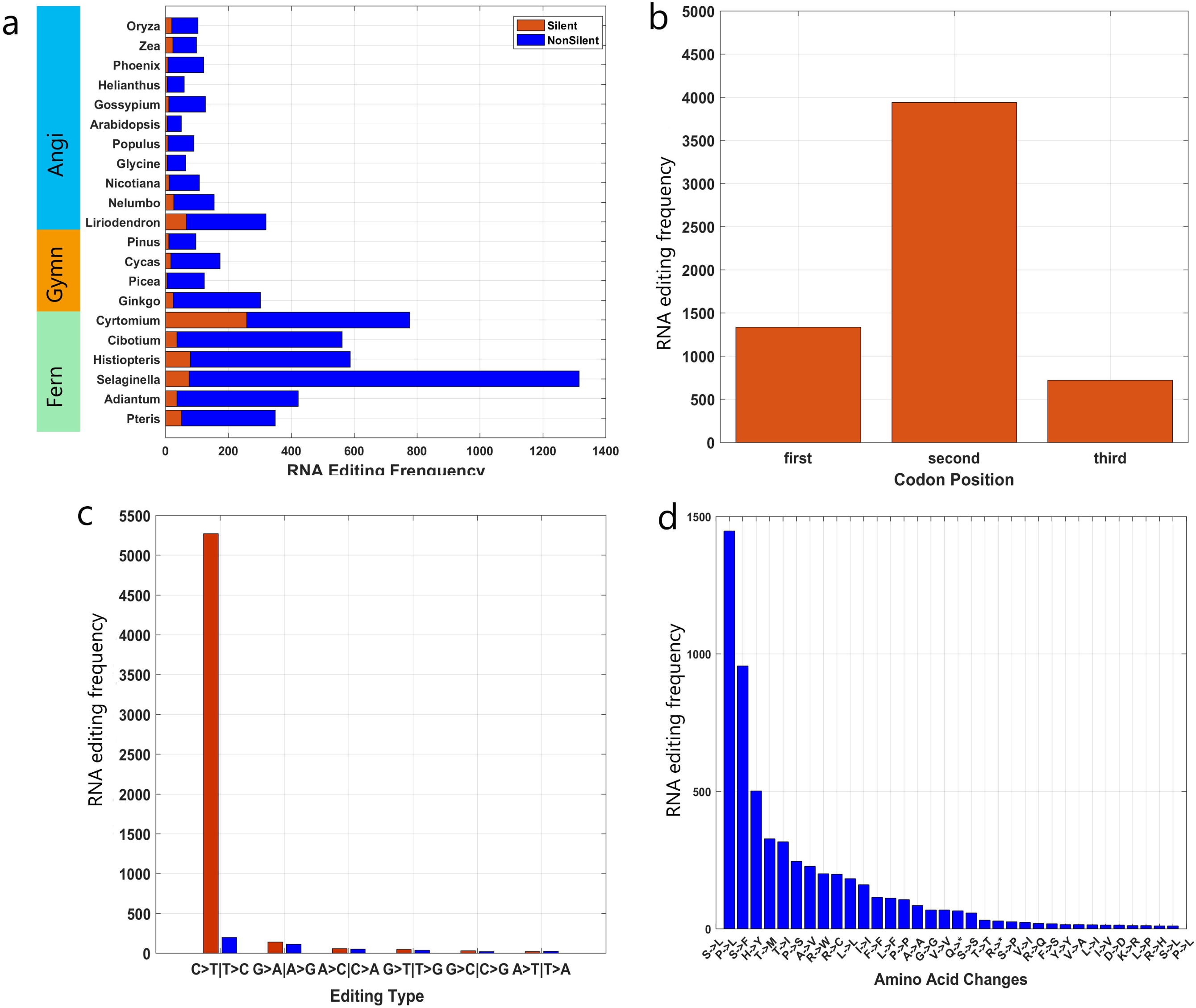
The statistics of identified RNA editing sites in chloroplast across 21 plants. (a) Total number of editing sites in all protein-coding genes across 17 angiosperms. Stacked bars depict numbers of nonSilent editing sites (blue), and silent editing sites (red) respectively. To simplify our presentation, the symbol of each species was represented by its first word of scientific name, such as Oryza -- *Oryza sativa*. (b) Codon position statistics of RNA editing sites. (c) Statistics of 12 RNA editing types, each pair was classified by two color bars (blue and red). (d) Statistics of amino acid change types.

All the filtered editing sites were located in 1,038 genes across all species, the average percent of edited genes in the three lineages (ferns, gymnosperms and angiosperms) are 0.85, 0.61, and 0.39 respectively (Table 1). Compared to the latter two lineages, more sites and genes were edited in chloroplast transcripts from ferns remarkably, see Figure 1a. In ferns, the largest number of editing sites was found in *Selaginella moellendorffii*, with 1,316 editing sites that is in good agreement with previous studies [23], chloroplast RNA editing in *Selaginella moellendorffii* is nearly 100-fold more abundant than in flowering plants, exclusively of the C-to-U type, involving 77 genes edited (∼ 96%), the overwhelming majority (94.3%) of the 1,241 silent editing events leaving codon altered directly. Differ from *Selaginella moellendorffii* that exclusively belonging to taxa of *lycopsida*, the other five fern plants are members of *Leptosporangiopsida*, has relatively smaller numbers of editing sites, represented by *Cyrtomium fortune*, owning second-largest number of editing sites among all specifies, with 610 editing sites and 79 edited genes (∼ 91%). Whereas for gymnosperms, the average number of editing sites and percent of edited genes were all less than that of ferns. Compared with other three species of the same taxa, *Ginkgo biloba* has the most editing sites, with 291 editing sites and 68 edited genes (∼81%) in chloroplast, reflecting its ancient nature of “living fossil”. On the opposite end, angiosperms have the lowest average numbers of editing sites and only a part of genes were effectively edited. It was noticeable that *Liriodendron tulipifera* and *Nelumbo nucifera* distinguished them from other angiosperms with more editing events. *Liriodendron tulipifera* is one of the most ancient flowering trees as the genus dates back about 65 million years, a total of 265 editing sites were detected, which is nearly 3-fold more abundant than that of other angiosperms and gymnosperms except *Ginkgo biloba*, the percent of edited genes was up to 75%, well above the average, reflecting early angiosperms possess much more diversified editing sites. The average number of editing sites among other 9 angiosperms showed no significant differences among them, but they were significantly lower than those of *Liriodendron tulipifera* and *Nelumbo nucifera.* The above results illustrated the differential distribution of RNA editing among varied plants, the cases of *Selaginella moellendorffii* and *Liriodendron tulipifera* also implies independent origins and subsequent evolutionary trajectories of editing processes.

### Variability of RNA editing among plants and genes

To further explore the evolutionary trajectory of RNA editing among different species with varied evolutionary relationships, for each gene, we summed the number of the editing sites across 21 plant species (see Additional file 4: Table S4), and picked out the top 30 genes with most editing sites across 21 plants for cluster analysis, as shown in Figure 2. We found that not every gene were edited in all the species, while a lack of editing at a few genes for certain species may be explained by two reasons, one is lack of the genes that annotated in chloroplast genome, another is no RNA editing occurred in the genes indeed. By grouping the genes based on their function, genes encoding membrane subunits of the chloroplast NDH complex and RNA polymerase exhibited the largest average numbers of editing sites, while ribosomal subunits showed the lowest numbers, this is consistent with previous studies in which RNA editing occurred preferentially in genes encoding membrane-bound proteins and genes under strong selection [24]. Due to the well-studied and abundant editing sites within the plant kingdom, *ndhB* gene is assumed to be a good example for the study of the conservation and evolution of RNA editing, in our study, *ndhB* was also confirmed to possess the most editing sites based transcriptome data, with 333 editing sites spread across 17 species. However, there is a biased distribution of RNA editing sites in *ndhB* among three linkages, in fern group, 50 sites were detected in *Selaginella moellendorffii*, and about 10 sites in the other ferns, in angiosperms, there were about 20 editing sites in each of the 11 angiosperms. In gymnosperms, RNA editing *ndhB* was only detected in *Ginkgo biloba*, for *Picea abies* and *Pinus massoniana*, no *ndhB* gene found in their chloroplast genome, whereas for *Cycas revolute*, no RNA editing events were detected in its *ndhB* gene, which may result from loss of editing or too low depth around genomic regions of its *ndhB* gene.

**Fig. 2.**
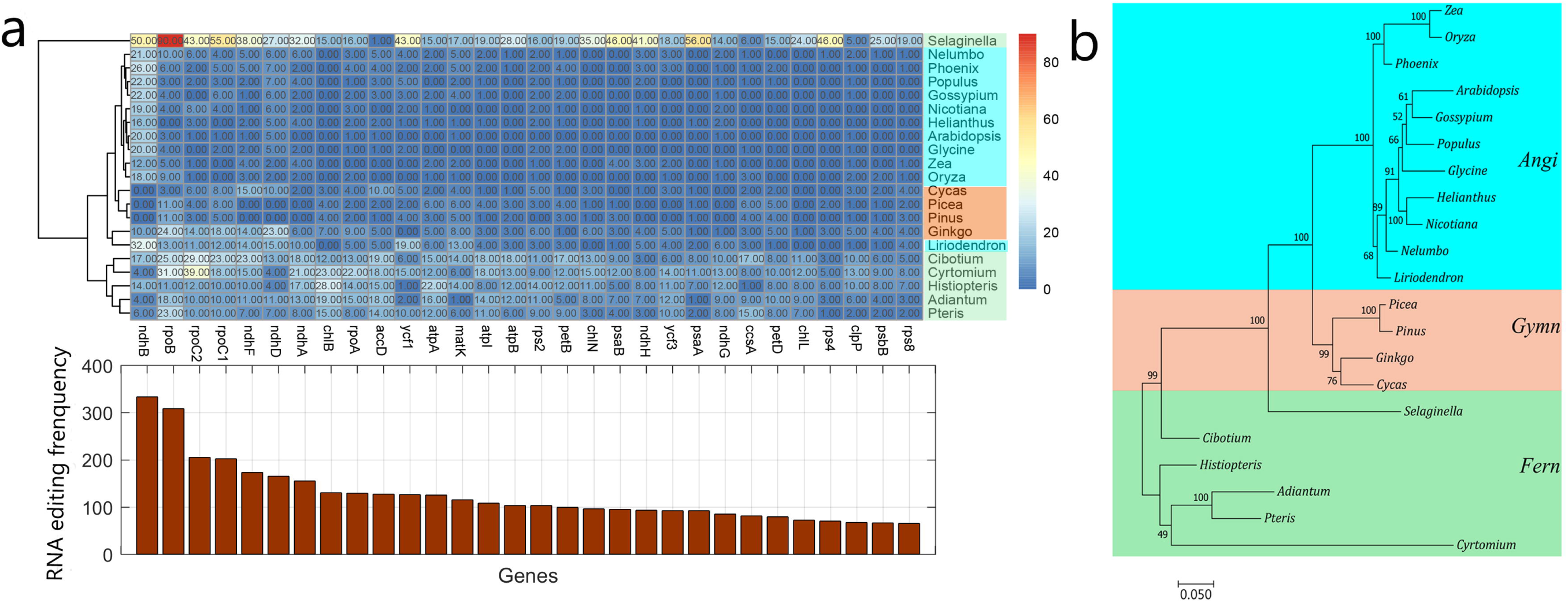
Analysis of numbers of RNA editing sites in top 30 genes with most editing sites across 21 plants. (a) Hierarchical cluster of numbers of RNA editing sites in top 30 genes was shown above, the x axis represents different genes, and the y axis represents plant, the total number of editing sites in each gene was shown below correspondly. (b) Phylogenetic tree by Maximum Likelihood method based on alignments of merged protein sequence for top 30 genes across 21 plants.

To compare the number of editing sites among the top-30 genes, a matrix of numbers of editing sites across 21 species was produced, and a hierarchically-clustered heatmap was plotted in Figure 2, which showed that 21 species were divided into three clustering groups with two exceptions, and the clustering relationships agreed with their phylogenetic tree based on sequence alignments roughly. *Selaginella moellendorffii* and *Liriodendron tulipifera* were clustered far away their own linkages respectively, further implies independent origins and subsequent evolutionary trajectories of editing processes. Chloroplast RNA editing may break out in early-branching plants in different linkages simultaneously and suffer a lot of loss during evolution. Hierarchically-clustered heatmaps of numbers of all RNA editing genes in chloroplast across 21 plants and each linkage were shown in Additional file 6: Figure S4-7 respectively.

### Uneven distribution of RNA editing extent

RNA editing extent was used to measure to what extent the edited transcripts among all transcriptome for one gene, if one site was edited, the C/G base (wild type) should be altered to the T/A base (edited type), thus its editing extent can be calculated by the formula: depth of edited bases (T and A)/total read depth of bases. In this study, we explored the distribution of editing extent among codon positions, species and edited genes. First, the comparison between codon positions showed that the distribution of RNA editing extent among them was uneven, and did not comply with the normal distribution, featuring an peak around ∼0.2 and fat tails, as shown in Figure 3. The average editing extent in second codon position (∼0.78) is higher than that of first (∼0.69) and third codon positions (∼0.66), suggesting non-synonymous substitution occurred in second codon position tend to be effectively edited, it was higher editing extent that dominated the landscape of RNA editing, and a selective advantage was given by a greater increase in hydrophobicity. Second, The average editing extent also demonstrated diversity of types across the 21 plants, ranging from 0.43 to 0.87 (see Figure 4a), *Selaginella moellendorffii* has the lowest editing extent (∼ 0.43), far below efficiencies of other species. In gymnosperms and angiosperms, *Ginkgo biloba* and *Nelumbo nucifera* had the lowest editing efficiencies (∼0.6) respectively, which seemed that more numbers of editing sites detected in these early-branching plants might negative affect their editing extent. However, as an ancient plant in angiosperms, *Liriodendron tulipifera was* one exception, with editing extent up to 0.81. The editing extent was also analyzed in each gene individually, we averaged the RNA editing extent among the top-30 genes across 21 plants and the result also demonstrated an uneven distribution, as shown in Figure 4b, *matK* gene has the lowest editing extent (∼0.5), oppositely, editing extent of *atpA* gene was the highest, up to 0.88.

**Fig. 3.**
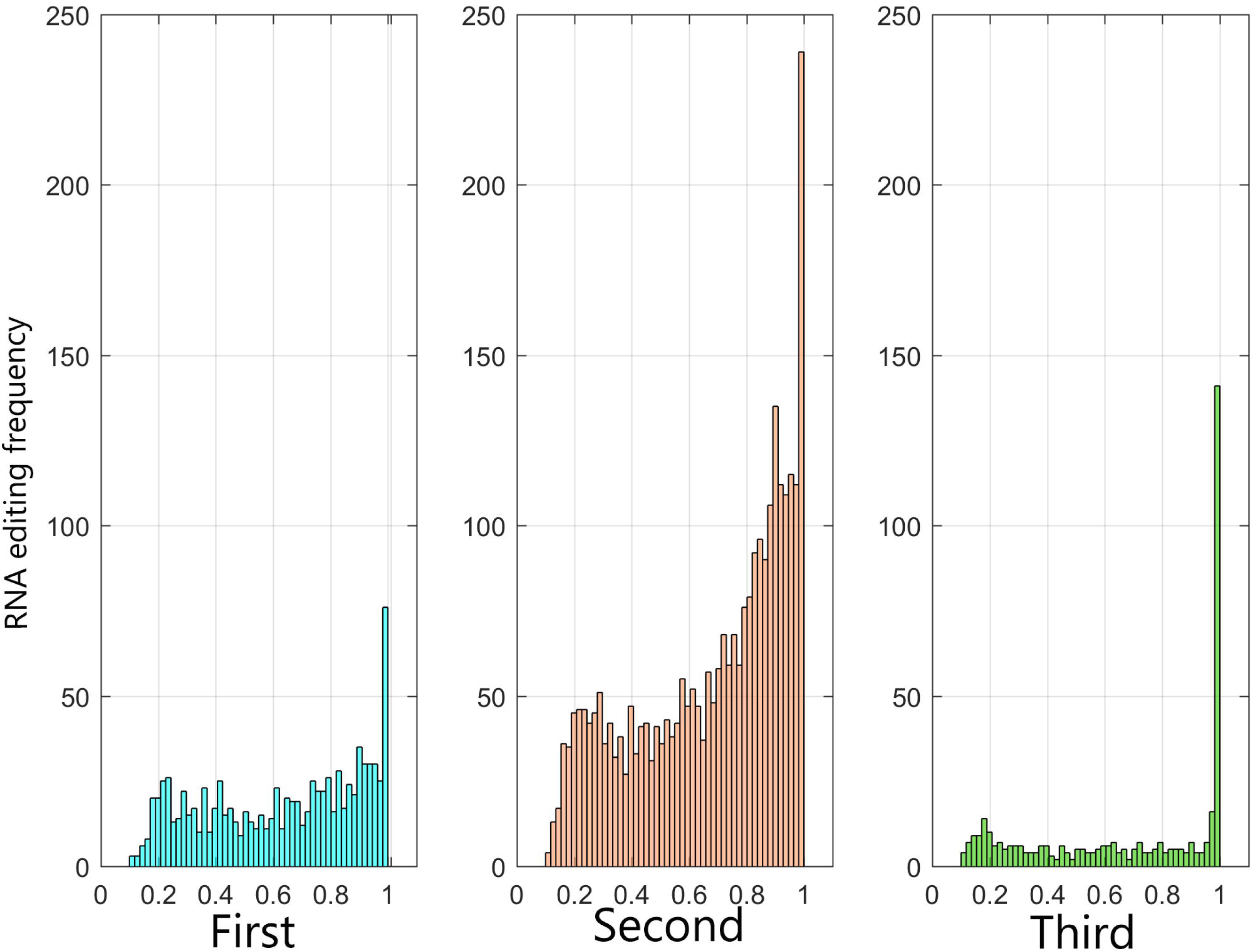
Editing extent of each identified editing site per codon position. The x axis represents editing extent, and the y axis represents frequency.

**Fig. 4.**
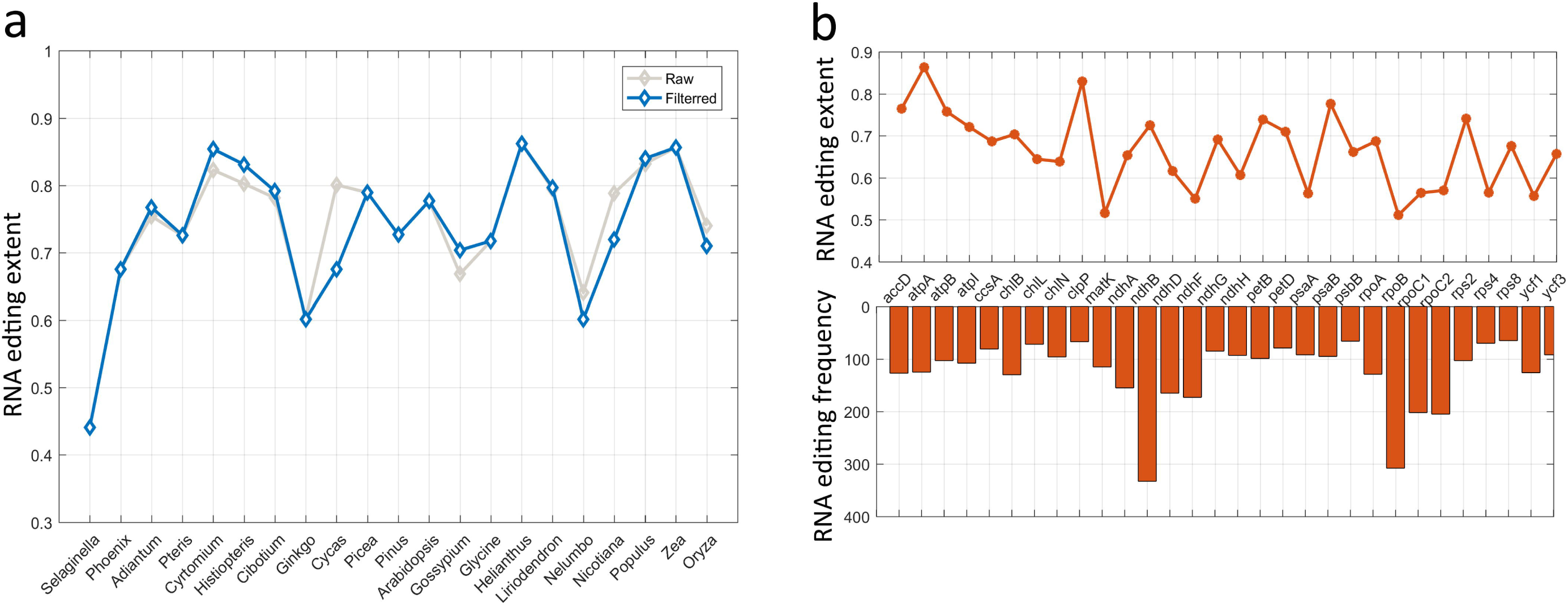
The statistics of identified RNA editing sites in chloroplast across 21 plants. (a) Distribution of average editing extent of all identified RNA editing sites in each plant specie. Grey and dark blue plots depict average editing extent for raw, filtered sites, respectively. (b) Distribution of average editing extent of top 30 genes. The above shows average editing extent of top 30 genes, the below shows numbers of RNA editing sites in top 30 genes correspondly.

### Reduced cytosines content with the evolution of plants

Considering the large differences in the scale of RNA editing events along with the evolution of plants, we analyzed the nucleic acid base content of involving genes shared by the 21 plants. There were a total of 51 genes annotated in all the chloroplast genome of 21 plants, for each plant, its corresponding gene sequences were extracted, the percent of cytosines for each gene was calculated by the formula: number of cytosines (C)/total number of bases (A/T/C/G), as listed in Additional file 5: Table S5. Afterwards, comparison between each two linkages were conducted, the cytosines content of each gene were averaged across all the members of each linkage, and two-tailed Wilcoxon rank-sum test was used to perform pairwise comparisons. The statistical result showed that a remarkable significance (p<0.05) was detected in the comparison of cytosines content between each two linkages except for gymnosperms-angiosperms, the percent of cytosines of ferns was far below than that of angiosperms, followed by gymnosperms, as shown in Figure 5a. The percent of cytosines of each gene across the 21 plants were further plotted in Figure 5b, which also demonstrated that the percent of cytosines dramatically declined roughly along with the evolution, with a few exceptions (such as *rpl23*). One striking example was *psbI* gene, has the highest percent of cytosines in *Selaginella moellendorffii* (∼0.37), and dropped to about 0.18 in other species. While the highest average of 51 shared genes was found in *Selaginella moellendorffii (∼0.26)*, the smallest average was found in *Glycine max* (∼0.17). Furthermore, *Ginkgo biloba* and *Liriodendron tulipifera* have higher average in angiosperms, correspond to ∼0.192 and ∼0.188 respectively, showed a positive correlation with their high numbers of RNA editing sites. The above results indicated the number of editing sites declined dramatically with the evolution of plants, which maybe due to loss of cytosine content in chloroplast gene for later-branch plants.

**Fig. 5.**
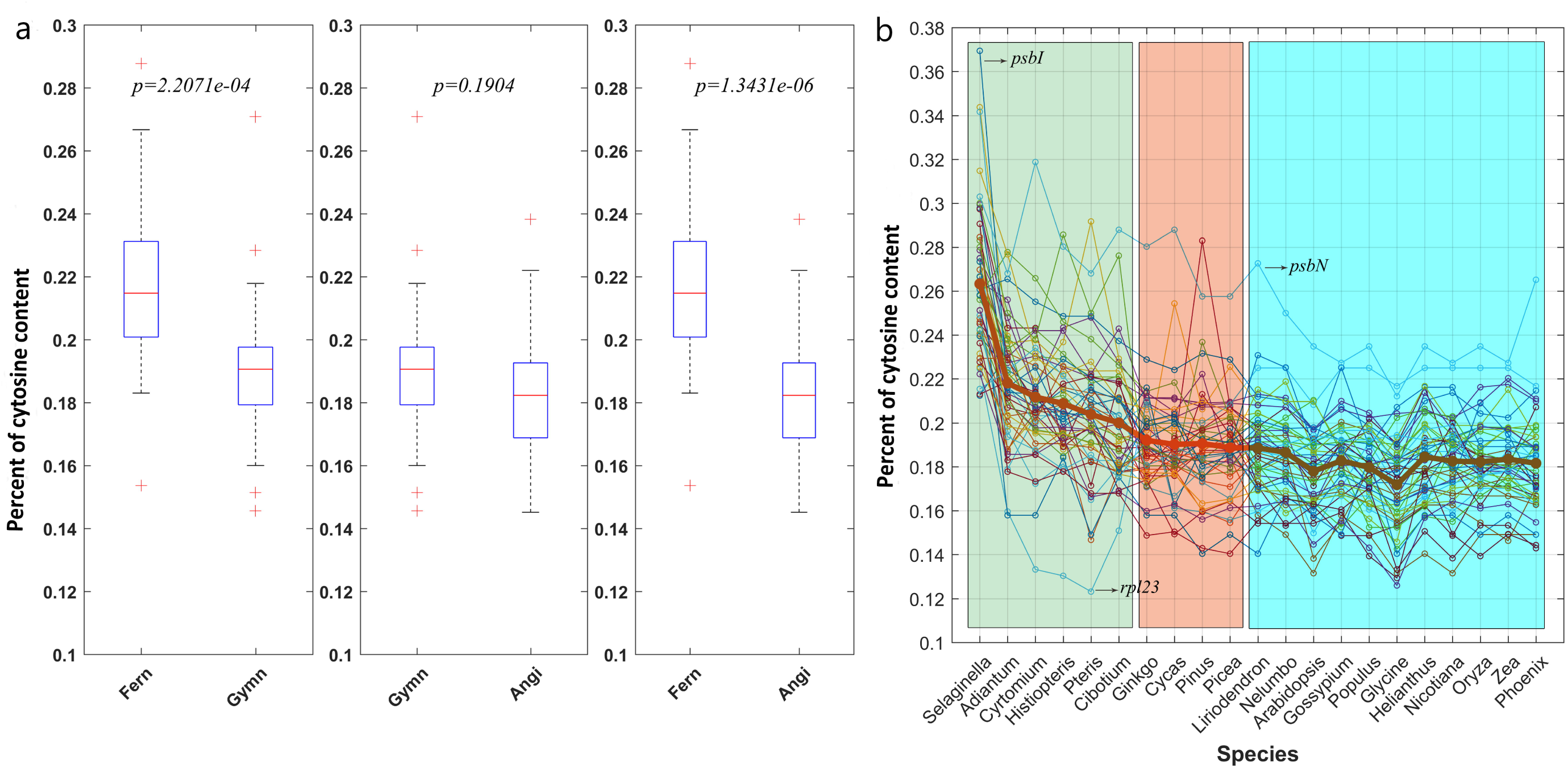
The statistics of cytosine content in shared RNA edited genes across 21 plant species. (a) Bar plots of pairwise comparisons between each two of groups (fern, gymn, and angi). Two-tailed Wilcoxon rank-sum test was used to perform the pairwise comparison. (b) Line plots of cytosine content for each shared RNA edited genes across 21 plant species. Average cytosine content of RNA edited genes for each specie is indicated by bold red lines. Two genes (*psbI* and *psbN*) are indicated by black arrows.

We illustrated one RNA editing example of *atpA* gene that may help to understand the evolution trajectory across plants vividly. The gene sequences of *atpA* across 21 species were collected, intersection of all the species’ RNA editing sites of *atpA* was concatenated for alignment and annotated in Figure 6. We marked all editing sites identified in this gene by yellow color, the distribution of its editing sites demonstrated that numbers of editing sites as well as cytosine content declined from ferns to angiosperms. We found that editing sites at third codon positions were poorly conserved, for example, in Figure 6a, RNA editing in site a was only occurred in *Adiantum aleuticum* in spite of existence of cytosine in other fern members, indicating that synonymous substitution were not active in the re-establishment of conserved amino acid residues in functional proteins. However, in contrast to RNA editing in site b that occurred in all three linkages, it is remarkable that RNA editing in site c was absent in gymnosperms, but occurred in several members of ferns and angiosperms. Whereas in site d, RNA editing occurred in two members of gymnosperms, and the base type of other 19 species were all thymine. The above results showed the diversity of RNA editing evolution, further validated that RNA editing in plant evolved independently in distant evolutionary lineages and new editing sites may occur occasionally in certain higher plants.

**Fig. 6.**
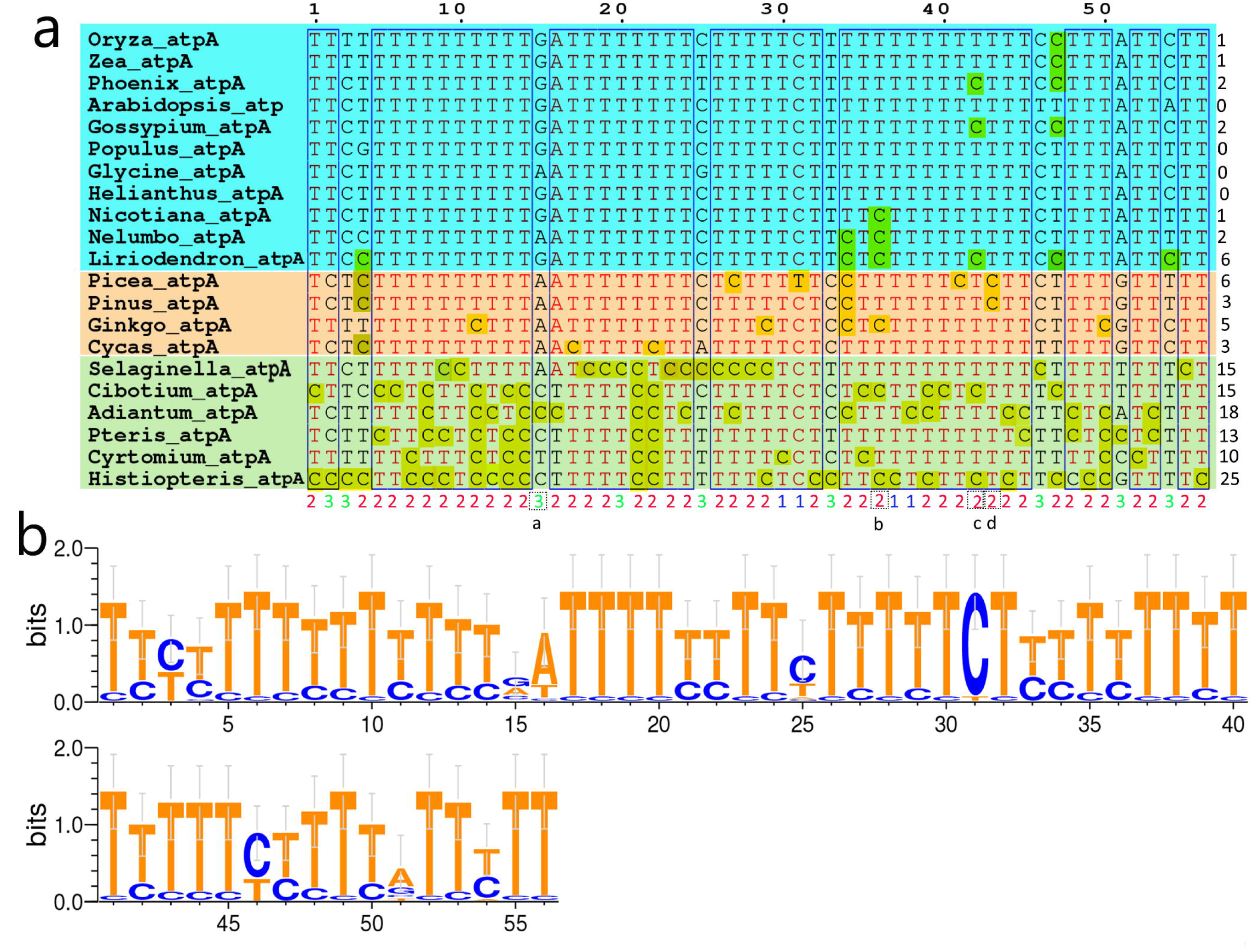
The statistics of cytosine content illustrated by RNA editing gene *atpA.* (a) Alignments of RNA editing sites extracted from initial RNA sequences in each specie, edited cytosines are marked by yellow. Codon positions are labeled by blue, red and green number under each column of site. (b) Sequence logo for RNA editing sites of *atpA* gene.

## Discussion

As a post-transcription process, RNA editing can modify the genome template to produce a different transcript [13]. Significant progress has been made in recent years in RNA editing studies for plant, numerous studies have proved that RNA editing occurred in nearly all plants in the kingdom, and demonstrated that RNA editing played roles not only in abiotic stress tolerance but also likely in the plant development, such as flower development and male sterile [8, 9, 14, 19], RNA editing may also result in secondary structure transformation of transcripts [25]. Until now, there are two viewpoints about the nature of RNA editing, one is contribution to proteomic sequence variation, thus provide another mechanism for modulating gene expression; another point thinks that RNA editing in plants is a repair mechanism to correct genomic point mutations at RNA level and alter the substitutional rate that is extremely low in organellar genes [26, 27].

With prior knowledge of detection of editing sites, primers are designed and RT-PCR products are performed to determine whether RNA editing occurs by comparing PCR products with the genomic DNA sequence. However, the prior approach is time-consuming and prone to underestimate editing sites and overestimate editing extent, because when comparing cDNA with genomic sequences for editing extent of less than 10%, one has to sequence more than 10 clones to find one edited transcript. With the advent of sequencing technology, the availability of large quantities of deep, strand-specific cDNA sequencing (RNA-seq) data offers a powerful technique to identify all the potential RNA editing sites and quantify their editing extent especially for the sites with very low extent, hence, RNA editing sites were identified in more and more organisms based on RNA deep sequencing [14, 19, 28].

In this study, to gain a better understanding of RNA editing in plant chloroplast, we collected a large mount of RNA-seq data and performed a series of bioinformatics procedures to investigate RNA editing in chloroplast across diverse plants that distributed in three plant linkages. A total of 5,389 editing sites located in leaf chloroplast genes across 21 plants were identified, many of the identified sites have not been previously reported and represent a valuable data set for future research community, demonstrating the powerfulness of RNA-seq data in identifying RNA editing sites. Agreed with previous study, the statics results showed RNA editing occurred in second codon position was mainly the largest, and majority (∼ 95%) of the editing events resulted in non-synonymous codon changes, additionally, a selective advantage was given by a greater increase in hydrophobic amino acid. We found that the cluster of numbers of RNA editing sites comply with the phylogenetic tree based on linked protein sequences approximately, further verified that the RNA editing across plant kingdom are comparatively conservative and accord with laws of evolution roughly. An uneven distribution of editing sites among species, genes, codon positions were found as well as the average RNA editing extent. In total, numbers of editing sites declined with the evolution of plants, *Selaginella moellendorffii* was identified to own the highest number of sites and the lowest editing extent (∼ 0.43), compared to other species. We also found that RNA-editing activities affecting third-codon positions showed a higher evolutionary variability. The number of editing sites declined dramatically with the evolution of plants, which maybe due to loss of cytosine content in chloroplast gene for later-branch plants.

Previous studies revealed that the evolutionary rate of organelle genomes is slower than that of nuclear genome, and result in accumulation of mutations of T-to-C, which might have been a prerequisite for the evolution of plant organellar RNA editing [29], RNA editing may have repaired numerous T-to-C mutations accumulating in the organelle genomes. During the evolution of land plants, many mutations would have been finally corrected to T in the genome, but at some sites, the residues may have been fixed to C to code for different amino acids. Our findings implied their independent origins across three linkages, chloroplast RNA editing may break out in early-branching plants in different linkages simultaneously, and suffer a lot of loss during evolution. The apparent reduction in the number of editing sites during the evolution of higher plants could indicate that RNA editing is disappearing in seed plants as genome mutations eliminate the need for editing at certain sites, since changes of genome are more constantly than RNA editing especially for certain functional proteins of great importance. The loss of editing sites along angiosperm evolution is mainly occurring by replacing editing sites with thymidines, hence, the increasing modification of C-to-T at the genome level might be more accurately to describe the evolution trajectory instead of loss of RNA editing sites. Those results suggest that the genome mutations is a widespread driving force underlying the loss of editing sites in angiosperm chloroplast.

## Conclusion

A systematic characterization and comparison of RNA editing events across major clades of plants could help to illuminate the evolution mechanism in a genome-scale patterns. Here, our goal was to document RNA editing in the chloroplast genome of 21species from three linkages and compare the distribution of editing sites among them. Based on a large amount of RNA-seq data, we used a relaxed automated approach combined with manual inspection to eliminate false positives, screened out thousands of RNA editing sites, demonstrating the advantages of combination of bioinformatics approach and sequencing data. Many of the identified sites have not been previously reported and represent a valuable data set for future research community. The analyses revealed that there is an uneven distribution of editing sites among species, genes, and codon positions, the average RNA editing extent also varied among different plant species as well as genes. The genome-wide distributions of chloroplast RNA editing across three linkages suggest that a major shift occurred on the lineage leading to seed plants, and plants have undergone drastic changes in both the numbers and patterns of editing. Our comparative study provided valuable information for evolution of RNA editing in plants. However, further mechanistic studies are still needed to characterize the enzymes and RNA-binding proteins involved in site recognition and editing across different clades.

## Methods

### Data collection

All data sets used in this study are publicly available. We selected 21 plants across three plant lineages (fern, gymnosperm, and angiosperm) for analysis of chloroplast RNA editing events. For each plant species, the corresponding transcriptome data was downloaded from SRA database at NCBI based on two criteria: first, to increase reliability of editing sites, SRA accessions with paired-end reads that possess higher mapping specificity were preferred, second, for the reason that more chloroplast mRNA in leaf were extracted and sequenced compared with other tissues, RNA-seq data obtained from leaves of wild type individuals were only selected. Besides, for each plant species, the reference file consisting of chloroplast genome sequences and corresponding gene annotation file in ‘tbl’ format were also downloaded from the GenBank database. Detailed information of SRA data and reference files used in our study were listed in Additional file 1: Table S1.

### Read mapping and SNP calling

In total, the identification process of RNA editing sites can be decomposed into three steps: first read alignment, second the SNP calling, and third detection of RNA editing sites. For each plant species, in order to increase sequencing depth, we merged all the SRA accessions from the same species into one sample. The quality control of paired-end Illumina sequencing data were evaluated first by NGSQCToolkit, low quality sequence data were filtered out (cutOffQualScore<20) [30], then transcriptome data from each plant species was aligned against its chloroplast reference by hisat2 software under default parameters [31]. Afterwards, the alignment results were sorted, removed duplicates, indexed, and sorted by using SAMtools [32]. Finally, the bcftools tool was used to identify SNPs, and VCF files that describe transcriptome variation were generated [33].

### Detection of RNA editing sites

The principles of RNA editing detection is similar to the transcriptome variation in SAMtools. Thus, we used the preliminary output from variant calling software to identify RNA editing sites. For each plant species, based on the SNP-calling results (in “VCF” format) and genome annotation files (in “tbl” format), the RNA editing sites were identified under default parameter values by using the REDO tool [21]. REDO is a comprehensive application tool for identifying RNA editing events in plant organelles based on variant call format files from RNA-sequencing data. REDO only works with three input files: the variant call format (VCF) files (records for all sites), the genome sequence file (FASTA format), and the gene annotation file (feature table file in “tbl” format, www.ncbi.nlm.nih.gov/projects/Sequin/table.html). Afterwards, regarding the high false positive of editing sites, REDO uses very stringent criterias to filter the raw variants by comprehensive rule dependent and statistical filters, as the below following: (1) quality control filter (MQ >255), the low quality sites are filtered according to the reads quality, (2) total reads depth filter (DP > 1), (3) alt proportion filter (alt proportion < 0.1), (4) multiple alt filter, only the variant with one alt allele is retained for RNA editing detection, (5) distance filter, the variant sites in short distance (<3 bp) are filtered out due to the possible positional interference for RNA editing, (6) spliced junction filter, variants within short spliced anchor (<2) are removed, (7) indel filter, the indel variants are removed, (8) likelihood ratio (LLR) test filter, LLR test is a probabilistic test incorporating error probability of bases (error probability is obtained using adjacent nonvariant sites in specific window) for detecting RNA editing sites (LLR <10), (9) Fisher’s exact test filter (p value < 0.01), the significance for a given RNA editing site (alt reads, ref reads) by comparing its expected levels (0, alt reads + ref reads) using the Fisher exact test, (10) complicated filter model, based on the statistics results for the attributes of codon table and experiment validated RNA editing sites, a complicated filter model was built according to five features of RNA editing sites, which are RNA editing types, alt proportion, amino acids change, codon phase, and hydrophobic/hydrophilic change. Finally, all raw RNA editing sites were detected, meanwhile, their corresponding annotation information files were also generated.

To minimize the number of false negatives with the automated approach, for the produced raw RNA editing sites, we manually examined all mismatches to examine the potential sources of error and eliminate false positives, and only kept the C-to-U and U-to-C editing types and exclude mismatches with other editing types, such as A-to-C, T-to-A, etc. In addition, to evaluate the reliability of editing sites number. Additionally, for each plant species, we also used PREPACT tools to predict potential RNA editing events using entire chloroplast genome as input file, with consensus prediction by at least 80% of the references under BLASTX mode, PREPACT originally relied on BLASTX hits in manually assembled collections of reference protein sequences [34].

### Characteristic statistics

All the filtered RNA editing sites detected in 21 plants were used for further statistics and feature analysis, including statistics of editing number, editing type, codon position, amino acid changes, involved genes and so on. In order to decipher the distribution of RNA editing frequency across different species, the top 30 genes with most editing sites across 21 plants were selected, cluster analysis and heatmap plotting were also provided based on the matrix of RNA editing numbers of top-30 genes across 21 plant species. The CDS sequences of top-30 genes across 21 plants were concatenated and subjected to alignments and phylogenetic tree construction using MEGA [35]. Meanwhile, the RNA editing extent of top-30 genes were also subjected to statistical analysis. In terms of RNA editing extent, its value at one site was expressed as the proportion between edited transcripts and total transcripts. If one site was edited, the C/G base (wild type) should be altered to the T/A base (edited type), since one editing site could be detected hundreds of times via sequencing, the number of wild type (C/G) or edited type (T/A) of bases could then be counted at this particular site, then the editing extent at one site could then be calculated by the formula: depth of edited bases (T and A)/total read depth of bases. Values of editing extent matrix were normalized by subtracting the row-wise mean from the values in each row of data and multiplying all values in each row of data by standard deviation value. For each lineage (fern, gymnosperm, and angiosperm), a heatmap was plotted across all of its species respectively using “pheatmap” function in R, the distance matrix of different samples was calculated using “dist” function with the default Euclidean method, and the hierarchical clustering was computed using “hclust” function.

Considering protein coding genes varied among different plant species, we picked out shared edited genes for the statistics of cytosine content across 21 plant species. For each shared edited genes, we extracted its CDS sequence, and calculated the ratio of cytosine content, and further performed pairwise comparisons between any two of lineages (fern, gymn, and angi). Two-tailed Wilcoxon rank-sum test was used. We illustrated *atpA* gene as a example, sequence logo of *atpA* gene was produced by WebLogo [36], alignment was constructed by using MEGA under default parameters [35].

## Additional files

**Additional file 1: Table S1.** SRA accessions and chloroplast genome for each plant used in the study.(XLSX)

**Additional file 2: Table S2.** Detailed information of filtered RNA editing sites across 21 plant species. (XLSX)

**Additional file 3: Table S3.** The statistical result of RNA editing sites in 21 plant species. (XLSX)

**Additional file 4: Table S4.** Numbers of RNA editing sites in 109 protein-coding genes across 21 plant species. (XLSX)

**Additional file 5: Table S5.** Cytosine content of RNA edited genes in 21 plant species. (XLSX)

**Additional file 6: Figure S1.** Average read depth of RNA-seq data across 21 species used in our study. **Figure S2.** Comparison of prediction for RNA editing sites between REDO and PREPACT3 tools. **Figure S3.** The attributes of RNA editing sites in chloroplast illustrated by samples of *Adiantum aleuticum*. **Figure S4.** Hierarchical cluster analysis of numbers of RNA editing sites in chloroplast across 21 plants. **Figure S5.** Hierarchical cluster analysis of numbers of RNA editing sites in chloroplast across 11 angiosperms plants. **Figure S6.** Hierarchical cluster analysis of numbers of RNA editing sites in chloroplast across 6 fern plants. **Figure S7.** Hierarchical cluster analysis of numbers of RNA editing sites in chloroplast across 4 gymnosperm plants. (DOCX)

## Abbreviations

C-to-U: Cytosine-to-uracil;
PPR: Pentatrico peptide repeat;
ndhB: NADH dehydrogenase subunit 2;
MORF: multiple organelle RNA editing factors

## Acknowledgments

We thank Yuepeng Han (Key Laboratory of Plant Germplasm Enhancement and Specialty Agriculture) for providing invaluable assistance in design of this study.

## Availability of supporting data

Supporting data are included as additional files.

## Authors’ contributions

ADZ, JF,XHJ, FPZ and XJZ conceived and designed the experiments, ADZ, JF performed data analysis, ADZ, and JF wrote the manuscript. XHJ, TFW and XJZ provided many critical suggestions. All authors reviewed the manuscript.

## Ethics approval and consent to participate

Not applicable.

## Consent for publication

Not applicable.

