## supplementary for "Gaining comprehensive biological insight into chloroplast RNA editing by performing a broad-spectrum RNA-seq analysis"

Wuhan 430072, China

**Figures:**

**Fig. S1** Average read depth of RNA-seq data across 21 species used in our study**.**

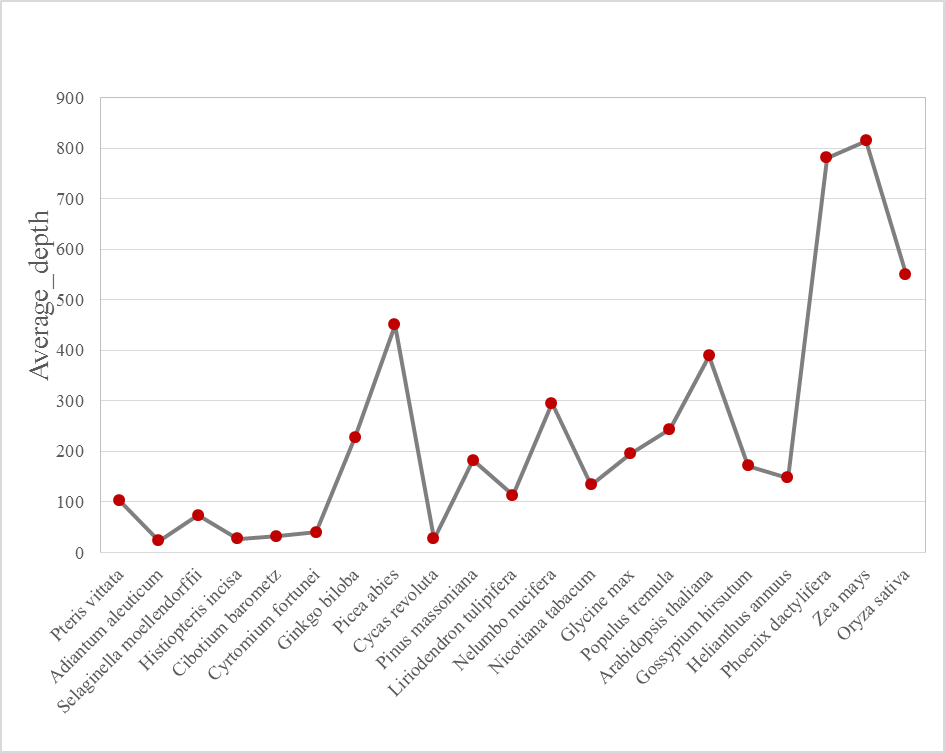

**Fig. S2** comparison of predicted numbers of RNA editing sites by RNA-seq data with website server.

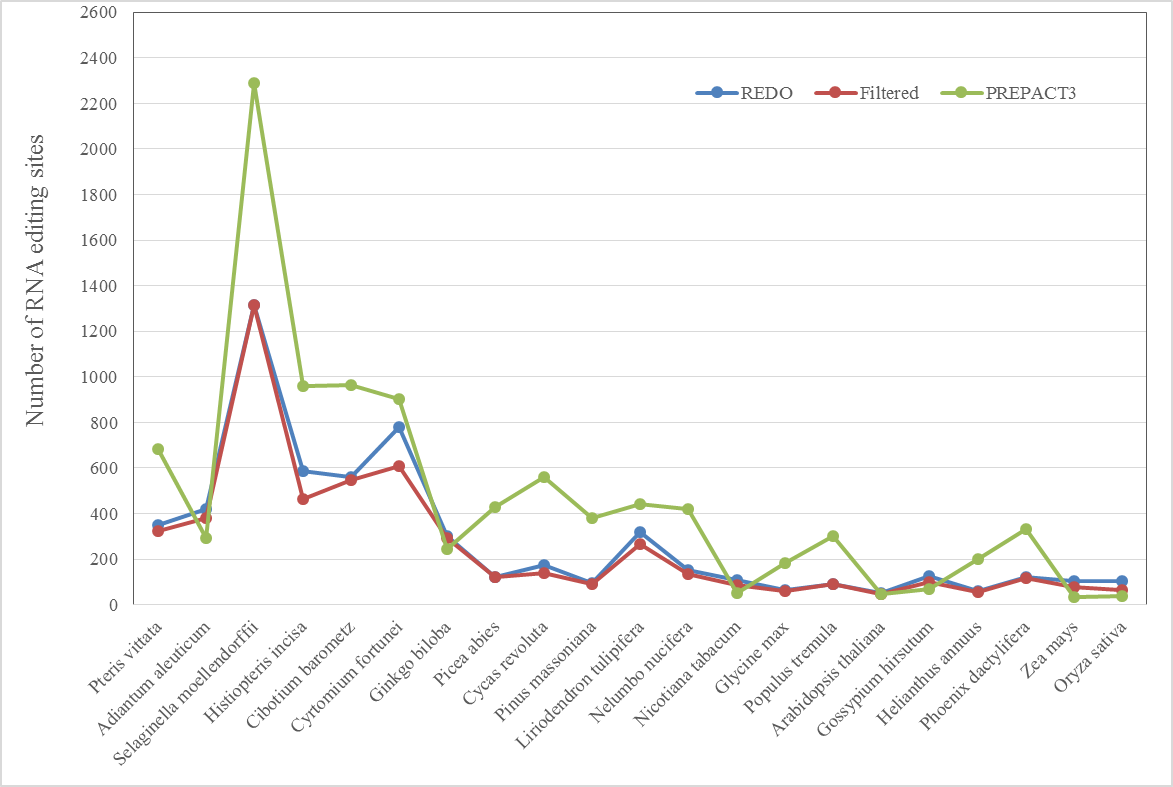

**Fig. S3** The attributes of RNA editing sites in chloroplast illustrated by samples of *Adiantum aleuticum*. The density of alternative allele proportion and sequencing error proportion, the density of total read depth and average read depth, the density of read proportion for alternative allele and reference allele, the density of distance to adjacent candidate editing site, the density of log10 LLR, the density of p value, the density of GC content, the boxplot of alternative allele proportion, the boxplot of sequencing error proportion, the boxplot of total read depth, the boxplot of total read depth/average read depth, the boxplot of distance to adjacent candidate editing site, the boxplot of log10 LLR, the boxplot of p value and the boxplot of GC content are shown for RNA editing sites; GC, GC content; LLR, likelihood ratio.

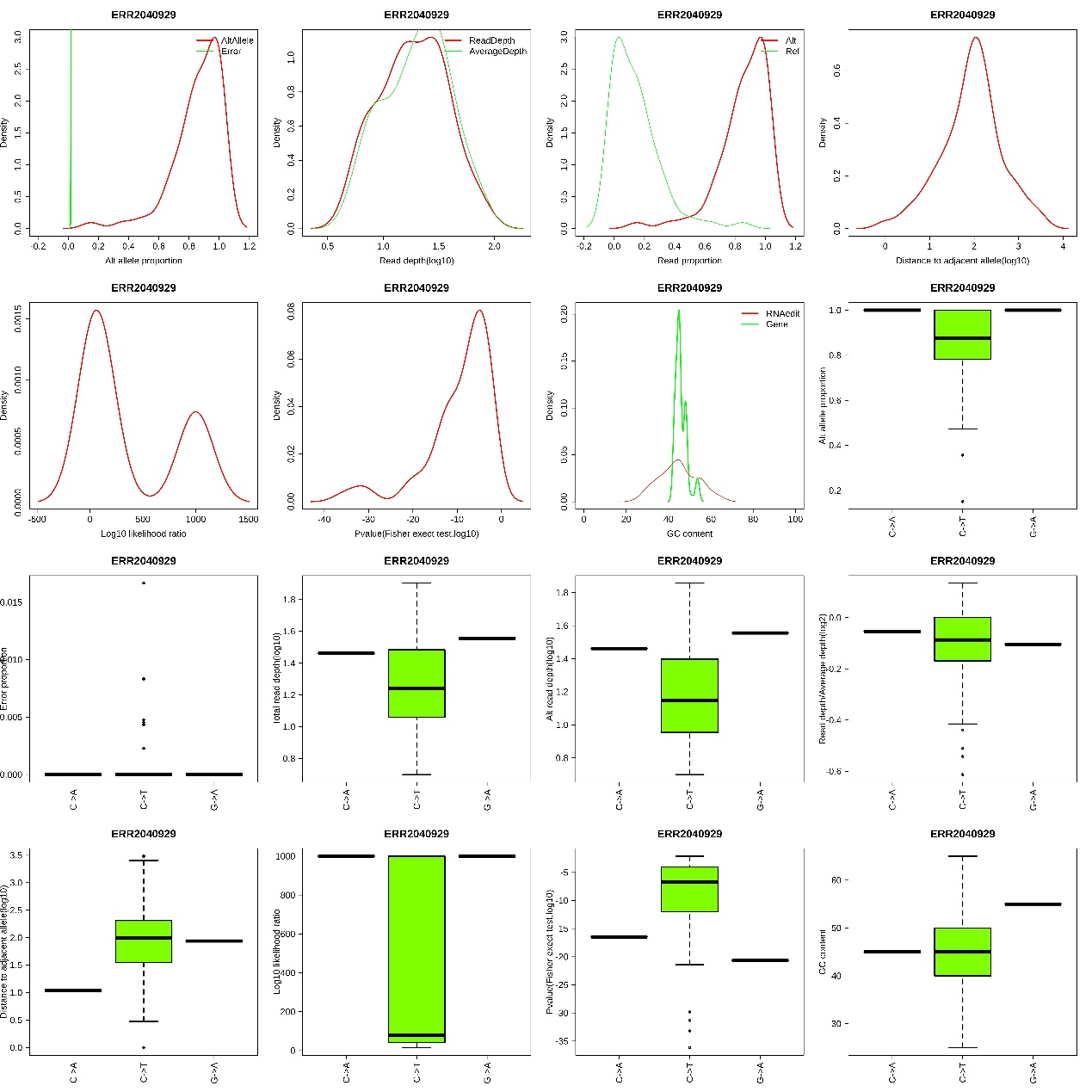

**Fig. S4** Hierarchical cluster analysis of numbers of RNA editing sites in chloroplast across 21 plants. The number of RNA editing sites for each gene was indicated inside the box. The x axis represents different plant species, and the y axis represents genes with RNA editing sites.

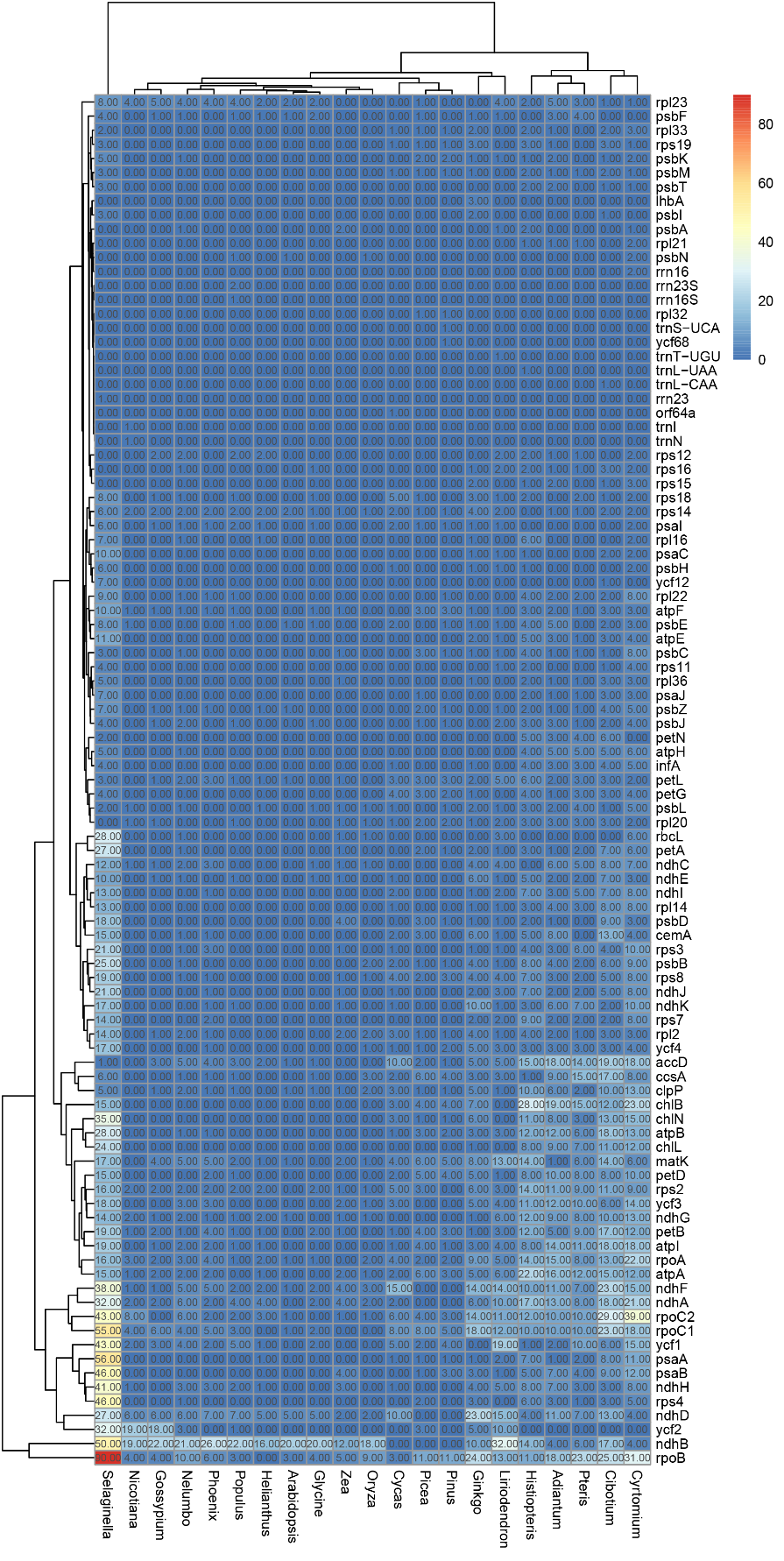

**Fig. S5** Hierarchical cluster analysis of numbers of RNA editing sites in chloroplast across 11 angiosperms plants. The number of RNA editing sites for each gene was indicated inside the box. The x axis represents different plant species, and the y axis represents genes with RNA editing sites.

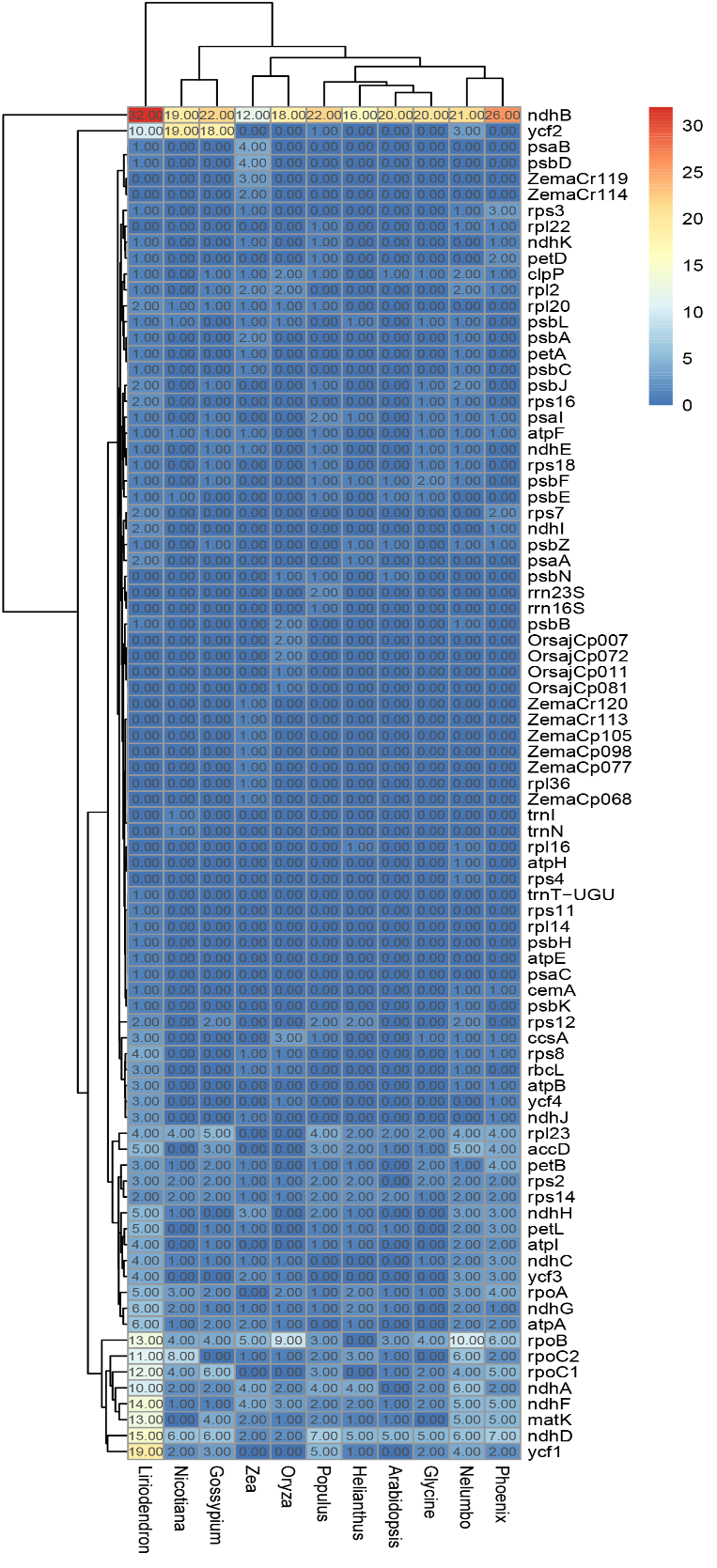

**Fig. S6** Hierarchical cluster analysis of numbers of RNA editing sites in chloroplast across 6 fern plants. The number of RNA editing sites for each gene was indicated inside the box. The x axis represents different plant species, and the y axis represents genes with RNA editing sites.

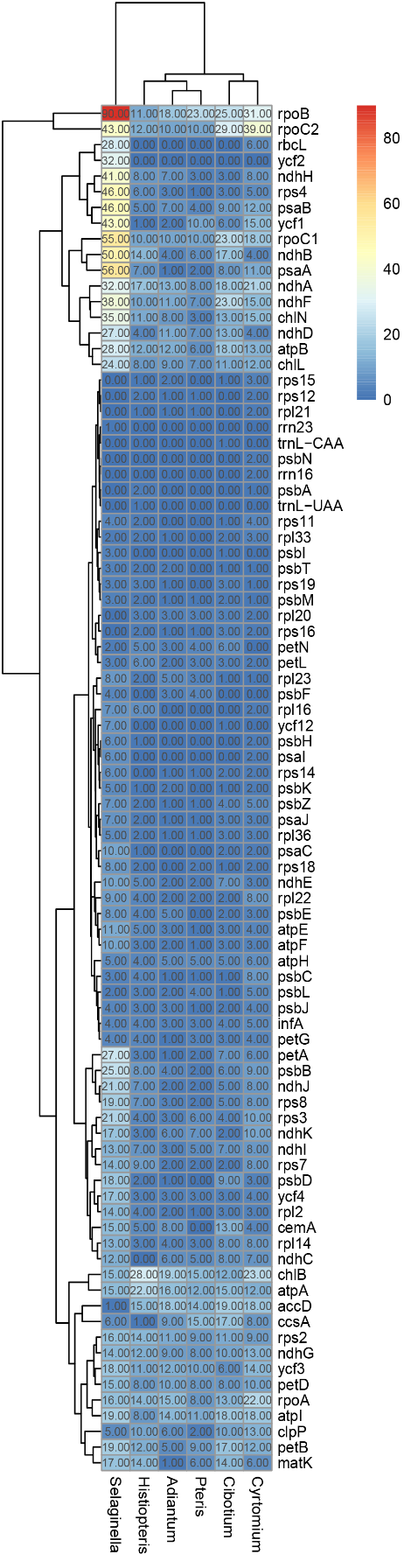

**Fig. S7** Hierarchical cluster analysis of numbers of RNA editing sites in chloroplast across 4 gymnosperm plants. The number of RNA editing sites for each gene was indicated inside the box. The x axis represents different plant species, and the y axis represents genes with RNA editing sites.

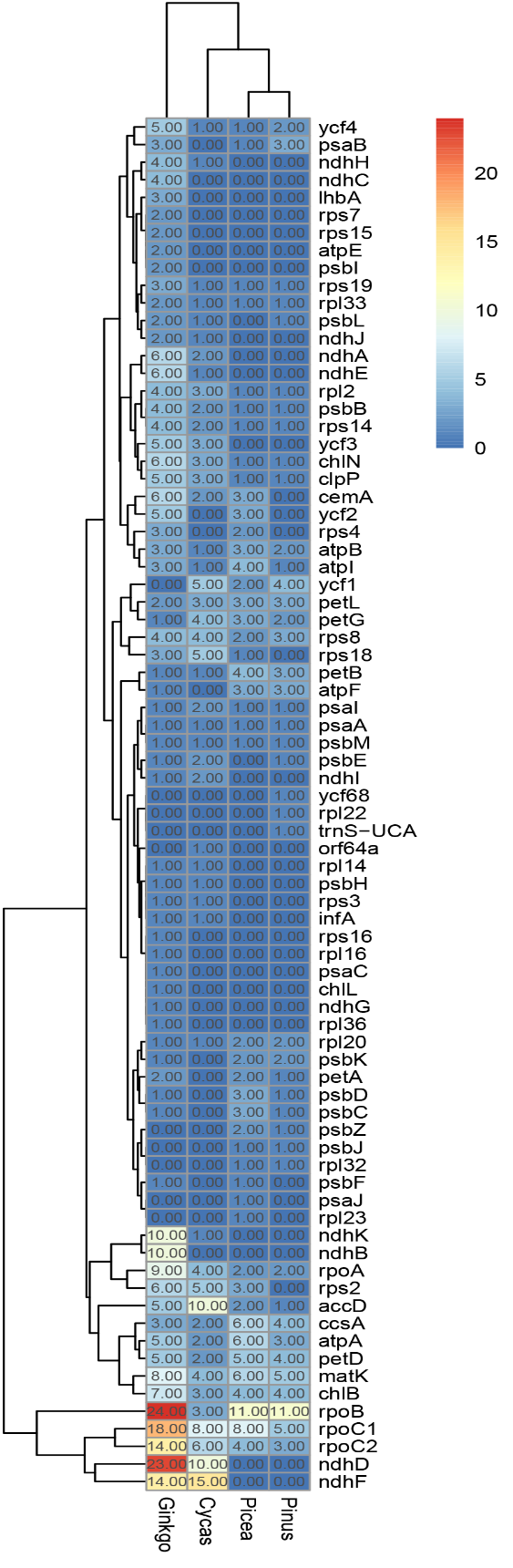
